# Identification of the lymphangioleiomyomatosis cell and its uterine origin

**DOI:** 10.1101/784199

**Authors:** Minzhe Guo, Jane J. Yu, Anne Karina Perl, Kathryn A. Wikenheiser-Brokamp, Matt Riccetti, Erik Y. Zhang, Parvathi Sudha, Mike Adam, Andrew Potter, Elizabeth J. Kopras, Krinio Giannikou, S Steven Potter, Sue Sherman, Stephen R. Hammes, David J. Kwiatkowski, Jeffrey A. Whitsett, Francis X. McCormack, Yan Xu

## Abstract

Lymphangioleiomyomatosis (LAM) is a metastasizing neoplasm of reproductive age women that causes cystic lung remodeling and progressive respiratory failure. The source of LAM cells that invade the lung and the reasons that LAM targets women have remained elusive. We employed single cell and single nuclei RNA sequencing on LAM lesions within explanted LAM lungs, known to contain smooth muscle like cells bearing mTOR activating mutations in TSC1 or TSC2, and identified a unique population of cells that were readily distinguished from those of endogenous lung cells. LAM^CORE^ cells shared closest transcriptomic similarity to normal uterus and neural crest. Immunofluorescence microscopy demonstrated the expression of LAM^CORE^ cell signature genes within LAM lesions in both lung and uterus. Serum aptamer proteomics and ELISA identified biomarkers predicted to be secreted by LAM^CORE^ cells. Single cell transcriptomics strongly supports a uterine neural crest origin of LAM^CORE^ cells; providing insights into disease pathogenesis and informing future treatment strategies for LAM.

**SIGNIFICANCE:** Present study identified a novel population of LAM^CORE^ cells, which is likely originated from uterine neural crest; identified novel LAM cell-specific secretome proteins that hold promise as potential biomarkers and therapeutic targets. Advancing the understanding of LAM pathogenesis and metastasis model may yield broader insights into the biology of cancer.

## INTRODUCTION

Lymphangioleiomyomatosis (LAM) is a rare, cystic lung disease primarily affecting women of childbearing age, which is associated with renal angiomyolipomas (AML), recurrent pneumothorax and progressive respiratory failure (Henske and McCormack, 2012). LAM occurs in patients with tuberous sclerosis complex (TSC), termed TSC-LAM, and also in women with a sporadic form of the disease (S-LAM), who do not have any evidence of heritable illness. In either case, LAM is caused by deleterious mutations in *TSC1* or *TSC2*, which encode components of a hetero-oligomer that controls the activity of the mammalian target of rapamycin (mTOR) (Smolarek et al., 1998). Cells within recurrent LAM lesions that occur in the allografts of some transplanted LAM patients arise from the recipient, supporting a metastatic mechanism for the disease (Carsillo et al., 2000). Mitotically quiescent cells within the LAM lesions (McCormack et al., 2012) are morphologically heterogeneous and include spindled cells with a smooth muscle phenotype, epithelioid cells that are variably positive for PMEL by HMB45 antibody staining, and stromal cells of mesenchymal, hematopoietic and endothelial lineages (Badri et al., 2013). LAM and AML share many morphological and immunohistochemical features, and are the founding members of the perivascular epithelioid cell neoplasm (PEComa) family of rare mesenchymal tumors that can arise in diverse tissues and exhibit pathological behaviors ranging from benign to malignant (Martignoni et al., 2008). The allele frequency of TSC1 or TSC2 mutations within LAM lesions range from 4% to 60%, with average less than 20% (Badri et al., 2013), supporting the concept that recruited stromal cells, rather than mutation-bearing LAM cells, dominate within the lesions (Badri et al., 2013; Clements et al., 2015; Johnson and Tattersfield, 1999). The mTOR inhibitor, Sirolimus, is an FDA-approved suppressive therapy that stabilizes lung function in most LAM patients (Gupta et al., 2019; McCormack et al., 2011); however, the drug does not eliminate LAM cells and can be associated with significant side-effects (McCormack et al., 2011). There is therefore an urgent need to develop new molecular targets and treatments for LAM.

Uncertainty regarding the origins of the LAM cells and the cellular and molecular mechanisms contributing to LAM cell migration, differentiation and dysfunction represent an obstacle to the development of new therapeutic strategies (Finlay, 2004). Although AMLs are cited as a potential source for LAM cells that invade the lung, not all LAM patients have these tumors (Avila et al., 2007; Henske, 2003). A neural crest cell origin has also been postulated (Delaney et al., 2014), in part because of the pluripotent, multi-lineage capacity of TSC mutant cells (which can differentiate into adipocytes or smooth muscle cells) and coexpression of melanocytic and smooth cell markers in the AML and LAM cells (Zhe and Schuger, 2004). Additional clues to LAM cell origins are derived from the remarkable gender restriction, anatomical distribution of disease and variation in disease progression across the female reproductive cycle. LAM is usually diagnosed in women of reproductive-age, suggesting an important role for female sex hormones in disease pathogenesis (Prizant and Hammes, 2016). Clinical observations that the menstrual cycle influences respiratory symptoms and occurrence of pneumothorax, and that decline of lung function slows after menopause and accelerates with exogenous estrogen use and pregnancy are consistent with this concept (Cohen et al., 2009; Gupta et al., 2019; Johnson and Tattersfield, 1999; Shen et al., 1987). Lymphadenopathy caused by LAM is more prominent in the pelvis and retroperitoneum than in the nodes in upper abdomen, thorax and mediastinum, suggesting a pelvic source for the neoplasm (Glasgow et al., 2008; Tobino et al., 2015). LAM lesions share some characteristics with uterine leiomyomas, including abnormal smooth muscle cells, expression of estrogen and progesterone receptors (Gao et al., 2014), and peak presentation during the reproductive years (Prizant and Hammes, 2016),. While pulmonary metastases from uterine leiomyoma are rare, some cases of benign metastasizing leiomyoma are associated with diffuse cystic changes and pneumothorax (Pacheco-Rodriguez et al., 2016), which supports the concept of lung restricted metastasis of morphologically quiescent uterine smooth muscle-like cells as a mechanism for cystic lung destruction. Eker rats with inactivating germ line mutations in *Tsc2* develop uterine tumors composed of estrogen responsive smooth muscle cells (Walker et al., 2003), and uterine specific deletion of *Tsc2* leads to estrogen-dependent myometrial tumors with smooth muscle and neural crest characteristics that spontaneously metastasize to the lung (Prizant et al., 2013; Prizant et al., 2016). Finally, a number of case reports and one small case series have described LAM lesions in the uteri of patients with S-LAM and TSC-LAM (Hayashi et al., 2011). Although indirect observations support the uterus as a potential source of cells that invade the lung, definitive proof in humans has been lacking, and this mechanism could not apply to the few cases of biopsy documented LAM in men that have appeared in the literature (Han et al., 2017).

Recent advances in single cell RNA-seq (scRNA-seq) enable detailed transcriptomic mapping of individual cells comprising organs and tissues, providing a unique opportunity to interrogate cellular heterogeneity, cell-cell interactions and cell-environment interactions in their native state (Ellsworth et al., 2017; Xu et al., 2016; Zhu et al., 2017). We utilized single cell RNA-sequencing, immunohistochemistry, and confocal microscopy to identify LAM^CORE^ cells and examine their effects on other cells in the LAM niche, to predict molecular and cellular network activity in LAM, to uncover LAM cell origins, and to identify useful biomarkers and molecular mechanisms underlying the pathogenesis of LAM.

## RESULTS

### scRNA-seq analysis of LAM lung explants identifies a unique cluster of LAM^CORE^ cells

Single cell and single nuclei RNA sequencing (scRNA-seq and snRNA-seq) were performed on LAM lung explants (LAM1/2/3/4; Figure 1; **Figure S1**), normal lung, renal AML, LAM uterine and normal uterine samples. scRNA-seq data of six additional female donor lungs from Gene Expression Omnibus (Reyfman et al., 2018) were included for integrative analyses. A total of 92,444 cells (21,964 cells from LAM/AML samples, 47,906 donor lung cells, and 22,574 cells from LAM/normal uterus samples) were used for further analysis (**Methods**). Clinical data, *TSC1* or *TSC2* mutations from LAM patients and control donors, and quality control statistics of single cell data are summarized in **Table S1**.

**Figure 1.**
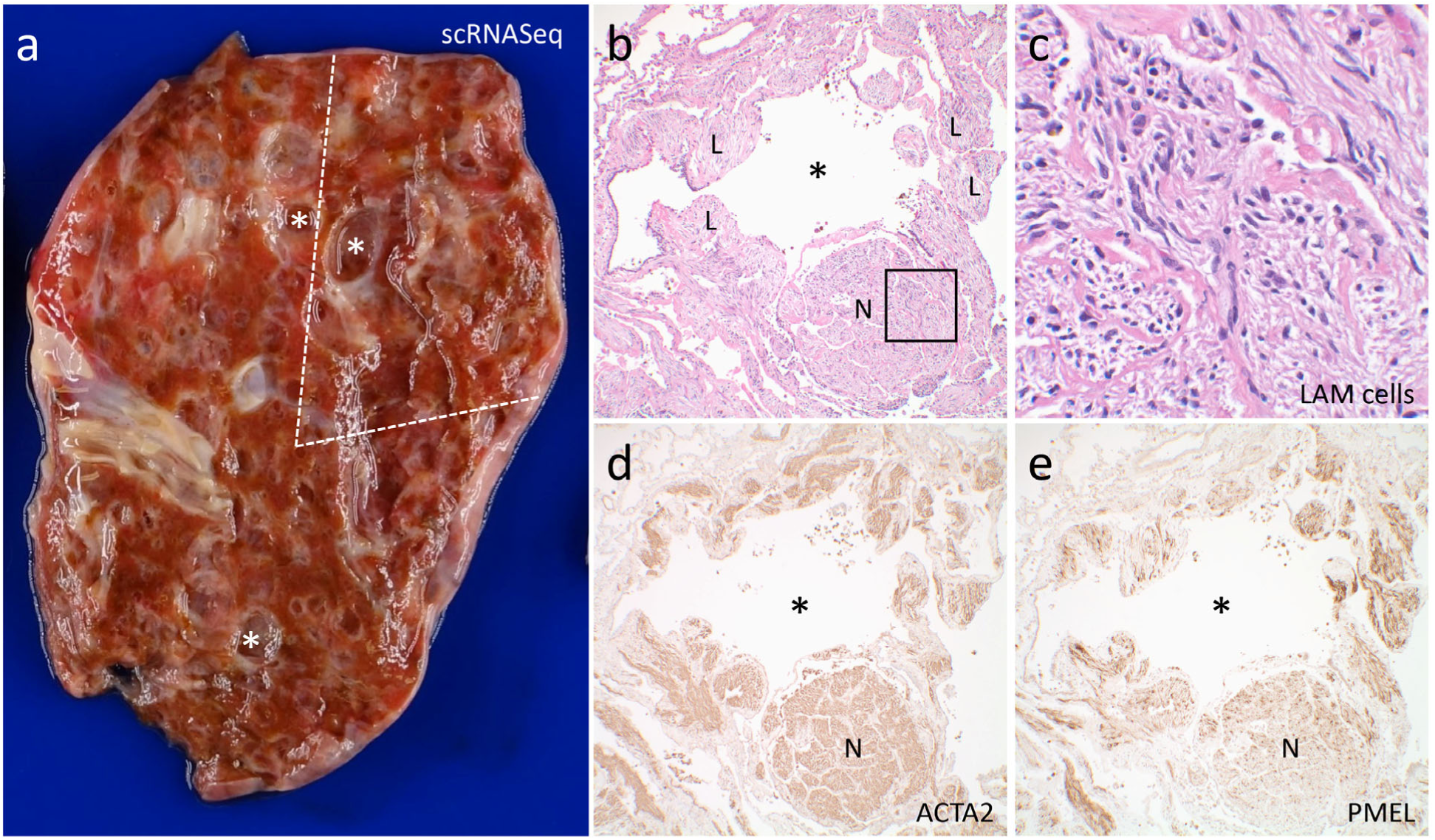
Pathology of representative LAM lung analyzed by scRNA-seq. **(a)** Fresh LAM lung (LAM3) with multiple variable sized cysts (*). The portion used for scRNA-seq analysis is indicated by dashed lines. **(b)** Histology of LAM3 lung adjacent to the portion used for scRNA-seq analysis showing a cyst (*) surrounded by LAM cells (L) arranged as bundles and as a nodule (N). **(c)** Higher magnification image of the boxed area in (b) showing haphazardly arranged spindled and epithelioid LAM cells with eosinophilic to clear cytoplasm. (**d-e**) LAM cells are diffusely positive for ACTA2 with a subpopulation also variably staining with the HMB-45 antibody for PMEL by immunohistochemical stain. Original magnification: 100x (b, d-e); 600x (c).

We identified 18 putative cell types from merged scRNA-seq data from LAM1 and LAM3 (Figure 2a; **Figure S2**). These included epithelial cells (alveolar type 1 (AT1), and alveolar type 2 (AT2) and airway (AirwayEpi)), vascular endothelial cells (vascular type 1(VasEndo-1) and vascular type 2 (VasEndo-2)), lymphatic endothelial cells (LymEndo), immune cells (T-cell, B-cell, macrophage, dendritic cell, monocyte, mast cell, natural killer cell (NK), and plasmacytoid dendritic cells), mesenchymal cells, and a small cluster of unique cells expressing multiple known LAM associated genes, including *PMEL* (Hoon et al., 1994), *ACTA2* (Zhe and Schuger, 2004), *ESR1* (Gao et al., 2014), and *FIGF (VEGFD)* (Seyama et al., 2006), hereafter termed LAM^CORE^ cells (Figure 2a, magenta; **Figure S2**). LAM^CORE^ cells were distinct from, but closely related to, an adjacent cluster of cells expressing typical lung mesenchymal cell signatures (Figure 2a, dark blue). Predicted cell types were visualized using cell type selective markers (Figure 2b; **Figure S2**).

**Figure 2.**
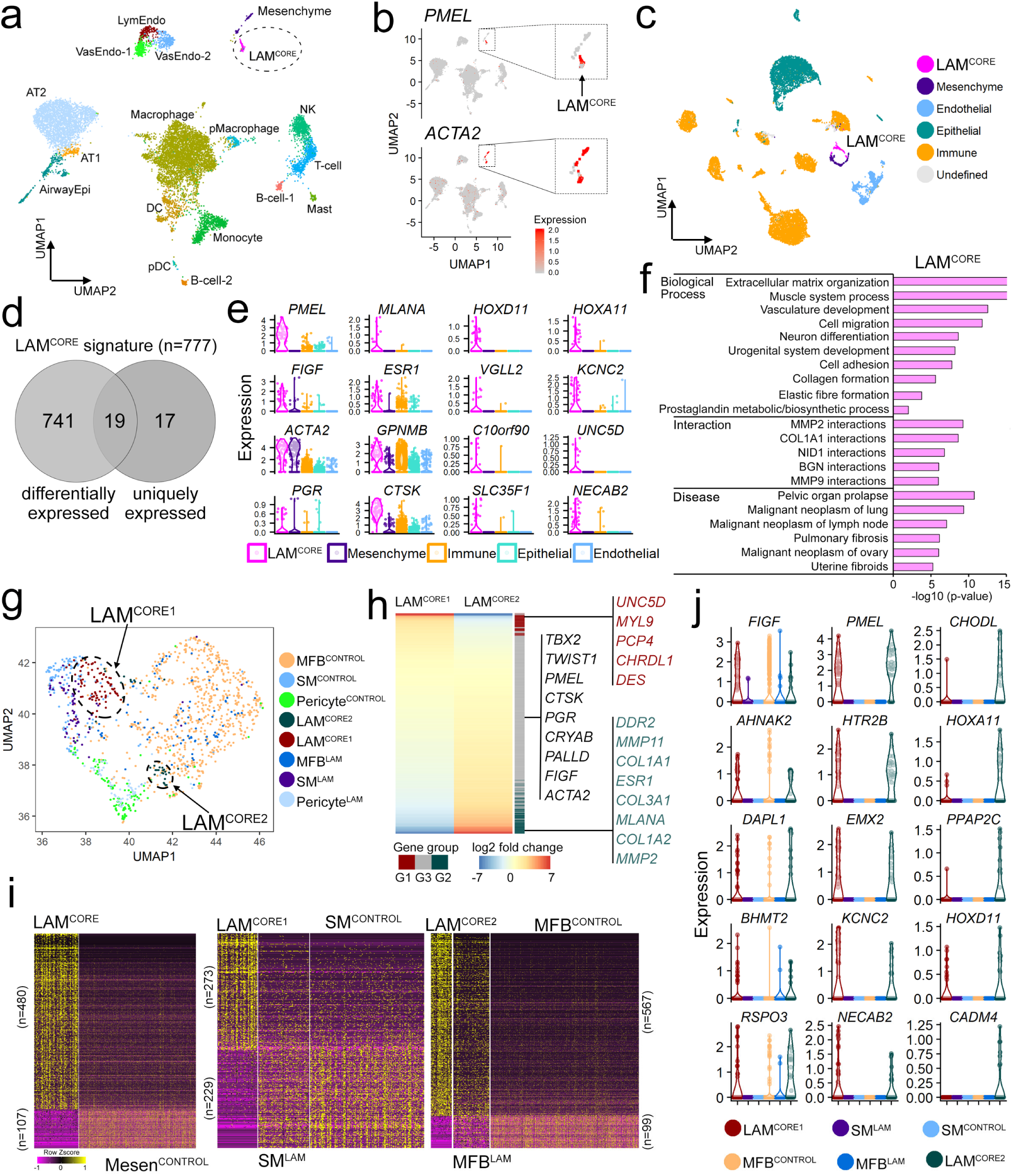
Identification of LAM^CORE^ cells and subtypes. **(a)** scRNA-seq data from LAM1 and LAM3. Dashed line encircles LAM^CORE^ cells that express known LAM markers. Cells are visualized by uniform manifold approximation and projection (UMAP) and colored by cell types. **(b)** Expression of known LAM markers in all cells of scRNA-seq data (left) and in LAM^CORE^ and LAM-associated mesenchymal cells in higher magnification (right). **(c)** Integrated scRNA-seq and snRNA-seq data from LAM lung tissues. Cells are visualized by UMAP and colored by major cell types. **(d)** The predicted signature for LAM^CORE^ cells is comprised of genes differentially expressed and uniquely expressed in LAM^CORE^ cells. **(e)** Violin plots represent expression of selective LAM^CORE^ signature genes in major cell types. scRNA-seq analysis identified both known LAM markers (left two columns) and novel markers (right two columns). **(f)** Functional enrichment analysis of the LAM^CORE^ signature genes. **(g)** scRNA-seq/snRNA-seq of LAM and control lungs identifies two subtypes of LAM^CORE^ cells (LAM^CORE1^, LAM^CORE2^), three subtypes of LAM-associated mesenchymal cells (SM^LAM^, MFB^LAM^, Pericyte^LAM^) and three subtypes of normal lung mesenchymal cells (MFB^CONTROL^, _SM_CONTROL_, Pericyte_CONTROL_)._ **(h)** Heatmap depicts differential expression of LAM^CORE^ signature genes in LAM^CORE1^ and LAM^CORE2^ subtypes. The analysis partitioned the LAM^CORE^ signature genes into three groups: “G1” genes differentially expressed in LAM^CORE1^ cells while “G2” genes in LAM^CORE2^ cells; “G3” genes were shared between the two subtypes. **(i)** Heatmaps of expression of genes differentially expressed in LAM^CORE^ vs. Mesen^CONTROL^ cells (left panel), in LAM^CORE1^ vs. SM^CONTROL^ (middle panel), and in LAM^CORE2^ vs. MFB^CONTROL^ (right panel). Mesen^CONTROL^ cells include MFB^CONTROL^ and SM^CONTROL^ cells. **(j)** Violin plots show genes selectively expressed in LAM^CORE1^, LAM^CORE2,^ or shared by both LAM^CORE^ subtypes.

scRNA-seq data from LAM1 and LAM3 were compared with single-nuclei RNA-seq (snRNA-seq) from LAM4 using the 10X Genomics platform (Figure 2c; **Figures S3-S4**). Interestingly, LAM^CORE^ cells were not detected in LAM2, perhaps related to ongoing use of Sirolimus in that patient. Consistent with the single cell data, no PMEL expression was detected by immunohistochemistry (IHC) in the smooth muscle LAM cells surrounding the cysts in the LAM2 lung despite positive PMEL expression in LAM1, LAM3, and LAM4 lungs (**Figure S1**), suggesting selective suppression of PMEL expression by mTOR inhibitor treatment. Since no LAM^CORE^ cells were detected in LAM2, data from this lung were not included in further analyses. Unbiased clustering of integrated LAM1/LAM3 scRNA-seq and LAM4 snRNA-seq data demonstrated concordance of all major cell types, including LAM^CORE^ and LAM associated mesenchymal cell clusters (Figure 2c; **Figure S4**). A total of 125 LAM^CORE^ cells were identified from the combined sc- and snRNA-seq analyses (Figure 2c).

Binomial and negative binomial probability-based tests (Shekhar et al., 2016) were used to identify genes differentially expressed in the cell clusters. The LAM^CORE^ signature consists of differentially expressed genes and uniquely expressed genes based on the criteria described in **Methods**. A total of 777 LAM^CORE^ signature genes were identified, including well known LAM markers, such as *PMEL, MLANA* (Busam and Jungbluth, 1999)*, FIGF, ESR1,* and many new markers, including *HOXD11*, *HOXA11*, *VGLL2*, *SLC35F1* (Figure 2d**-e**; **Table S2**). LAM^CORE^ signature genes were subjected to functional enrichment analysis and Ingenuity Pathway Analysis (IPA). As shown in Figure 2f, LAM^CORE^ signature genes were enriched in bioprocesses involving extracellular matrix organization, collagen and muscle fiber formation, cell migration and adhesion, as well as prostaglandin metabolic/biosynthetic process and development and differentiation of vessels, neurons, and the urogenital system. When compared with curated DisGeNET (a publicly available collection of genes and variants associated with human disease), LAM^CORE^ signature genes share most identity with disease genes associated with neoplasms and the genitourinary tract, including pelvic organ prolapse, neoplasm of ovary, uterine fibroids, and secondary malignant neoplasm of lung and lymph node (Figure 2f). Pathway analysis predicted the activation and interactions among mTOR, WNT (Mak et al., 2005), HIPPO (Liang and Pende, 2015), VEGF, PI3K-AKT, prostaglandin (Li et al., 2014), Rho GTPases, ILK, and ERK/MAPK (Gu et al., 2013) signaling pathways (**Figure S4**). While activation of mTOR, PI3K-AKT, ERK/MAPK pathways are known features of LAM, mapping them to stromal or epithelial tissue compartments has been historically challenging. scRNA-seq technology enables the linkage of these pathways to specific cell populations (i.e., LAM^CORE^ cells) within the diseased lung (**Figure S4**) in a manner that has not been previously possible.

### Single cell RNA-seq analysis identifies distinct pulmonary LAM^CORE^ cell subtypes

To understand and assess the heterogeneity of the LAM^CORE^ cell cluster, we compared the transcriptome of LAM^CORE^ cells with LAM-associated mesenchymal cells and mesenchymal cells from control female lungs (Reyfman et al., 2018). Unbiased graph-based clustering identified 8 mesenchymal cell subtypes, including two distinct LAM^CORE^ subtypes (LAM^CORE1^ and LAM^CORE2^), three LAM-associated mesenchymal cell subtypes: smooth muscle-like (SM^LAM^), matrix fibroblast-like (MFB^LAM^) and pericyte-like (Pericyte^LAM^), and three normal lung mesenchymal cell subtypes: SM^CONTROL^, MFB^CONTROL^, and Pericyte^CONTROL^ (Figure 2g; **Figures S5-S6**). LAM^CORE1^ cells expressed smooth muscle (SM) selective cell markers *ACTA2*, *ACTG2* and *DES*, and clustered most closely with SM^CONTROL^ and SM^LAM^ cells. LAM^CORE2^ cells expressed higher levels of matrix fibroblast (MFB) cell markers *TCF21*, *COL1A1* and *PDGFRA* and clustered closely with MFB^LAM^ and MFB^CONTROL^ cells, Pericyte^LAM^ co-clustered with Pericyte^CONTROL^ cells and expressed higher levels of *PDGFRB*, *NOTCH3, RGS5,* and *ACTA2* (**Figure S6**). Proliferation markers, including *MKI67*, *TOP2A*, *HMGA2* (D’Armiento et al., 2007), *BUB1* and *FOXM1* were not increased in LAM^CORE^ cell subtypes (**Figure S6**). LAM^CORE^ signature genes (n=777) were separated into three groups based on their expression patterns (“G1”, “G2”, and “G3” in Figure 2h) using the negative binomial probability based differential expression test (Yu et al., 2013) (“G1” genes for LAM^CORE1^, “G2” genes for LAM^CORE2^ and “G3” genes are shared between LAM subtypes) (Figure 2h; **Table S3**). Signature genes selectively expressed in LAM^CORE1^ cells included *UNC5D*, *MYL9*, *PCP4*, *CHRDL1* and *DES*. Signature genes selectively expressed in LAM^CORE2^ cells included *DDR2* (Ferri et al., 2004)*, MMP2/11* (Matsui et al., 2000), *COL1A1/1A2/3A1*, *ESR1* and *MLANA*. Shared signature genes included *TBX2*, *TWIST1*, *PMEL*, *CTSK* (Chilosi et al., 2009), *CRYAB* (Iwaki and Tateishi, 1991) and *PGR* (Gao et al., 2014) (Figure 2h).

Angiomyolipoma (AML) are present in approximately 30% of sporadic LAM and 70% of TSC-LAM patients (Henske and McCormack, 2012). AMLs consist of mesenchymal cells sharing morphologic characteristics and cellular markers with pulmonary LAM cells, and have been considered a potential source of LAM cell metastases (Henske, 2003). We compared scRNA-seq data from LAM^CORE^ cells with scRNA-seq from a sporadic renal AML. Hierarchical clustering of cells from pulmonary LAM and renal AML samples identified similar major mesenchymal cell types, i.e., smooth muscle like, matrix fibroblast like and pericyte like cells, **Figure S7**, consistent with the cells identified using the single cell data from LAM lung samples alone. Taken together, renal AML shares transcriptomic similarity with pulmonary LAM sharing both cell subtype identity and common signature genes.

### Distinction of LAM^CORE^ from normal lung mesenchymal cells

Genes differentially expressed in LAM^CORE^ cells and lung mesenchymal cells from control female lungs were identified by the negative binomial probability-based tests (Shekhar et al., 2016) (Figure 2i, left panel; **Table S4**). Bioprocesses and pathways induced in LAM^CORE^ cells compared to normal lung mesenchymal cells include the MAPK signaling pathway, cytoskeletal regulation by RHO GTPase, and FoxO signaling as shown in **Figure S8**.

Transcriptional changes distinguishing LAM^CORE^ subtypes and corresponding normal lung mesenchymal subtypes were identified by comparing LAM^CORE1^, which exhibited a smooth muscle gene signature, with smooth muscle cells from control lungs. LAM^CORE2^ cells, which exhibited a matrix fibroblast gene signature, were compared with control lung matrix fibroblasts. Differentially expressed genes in LAM^CORE^ subtypes and corresponding mesenchymal cell subtypes from control lungs are shown (Figure 2i, right two panels; **Table S5**). Genes that were uniquely expressed or shared by LAM^CORE^ subtypes, but distinct from normal lung mesenchymal cells were identified (Figure 2j). LAM^CORE1^ cell-selective markers include *FIGF*, *RSPO3* (R-Spondin 3) and *AHNAK2* (AHNAK nucleoprotein 2). *HOXA11*, *HOXD11*, and *CADM4* were among LAM^CORE2^ cell-selective markers. Genes expressed in both LAM^CORE1^ and LAM^CORE2^ subtypes and were distinct from control lung mesenchymal cells including *PMEL*, *NECAB2*, *HTR2B*, *EMX2* and *KCNC2* (Figure 2j). Gene Ontology enrichment analysis of signature genes of LAM^CORE^ subtypes identified subtype specific and shared bioprocesses and pathways that were induced in LAM^CORE^ subtypes compared to normal lung. Genes involved in the synthesis and degradation of glycosaminoglycans, which are referred to as ECM-resident growth factors (Nikitovic et al., 2014), were highly enriched in both LAM^CORE^ subtypes. Pathways and bioprocesses associated with cellular response to growth factors, to MAPK and NFKB signaling, were mostly increased in LAM^CORE1^, while those associated with cellular response to estradiol, Wnt, and RHO GTPases signaling were selectively increased in the LAM^CORE2^ cells (**Figure S8**).

### Transcriptomic changes in LAM associated lung epithelial and lymphatic endothelial cells

Complex changes in transcriptomic programs occur in multiple cell types, including lymphatic endothelial, airway and alveolar epithelial cells, and inflammatory cells in LAM (Osterburg et al., 2016). Lymphatic endothelial cells and associated markers (*LYVE1*, *PROX1*, *PDPN* and *CCL21*) were increased in LAM tissue (Davis et al., 2013). Expression of genes associated with invasion of lymphatic vessels during metastasis (*UNC5B*, *NECTIN2*, *CD200*, *ESAM*, *ENG*) (Clasper et al., 2008) was increased in LAM associated lymphatic endothelial cells (**Figure S9a-b**). *VEGFR3 (FLT4)* (Kumasaka et al., 2004), a known receptor of *VEGFD (FIGF),* was induced in LAM endothelial cells (**Figure S9c**). *VEGFC* was selectively expressed in a subset of LAM vascular endothelial cells (**Figure S9d**). Epithelial cell gene expression in LAM lung was distinct from that of controls. For example, expression of AT2 cell signature genes involved in host defense and innate immune response, sphingolipid biosynthetic process, and cytokine/interleukin signaling were increased, while several characteristic bioprocesses of normal AT2 cells, including biosynthesis and metabolism of lipids and lipoproteins, SREBP mediated cholesterol biosynthesis, and activity of the ERK1 and ERK2 cascade were suppressed in LAM associated AT2 cells. **Figure S9e** depicts representative genes (*FGF7, DMBT1, and LCN2*) induced in LAM AT2 compared to normal AT2 cells. The pleiotropic changes in gene expression in multiple cell types in the LAM lesions are consistent with the complexity of histopathological changes associated with pulmonary LAM lesions.

### Experimental validation of LAM^CORE^ cells types, signature genes, and pathways

In normal human lung, peribronchiolar and vascular smooth muscle cells stained for ACTA2 but not for PMEL (**Figure S10a-c**). In LAM lungs, LAM cells were diffusely positive for ACTA2, with a subpopulation of LAM cells co-expressing PMEL and ACTA2, and many fewer LAM cells expressing MLANA (Figure 1; Figure 2e**; Figure S1k; Figure S10d-f**). The increased expression of representative known LAM^CORE^ signature RNAs (*CTSK*, *FIGF*, *PGR*, and *MYH11*) and novel LAM^CORE^ signatures (*KCNAB1* and *PALLD*) were validated by qPCR and immunofluorescence staining (**Figure S10g-h**). LAM^CORE^ signature genes RAMP1 and MYH11 were co-expressed with ACTA2 in LAM nodules (Figure 3h-i**; Figure S10i**), validating RAMP1 as a new marker for LAM^CORE^ cells. In contrast, RAMP1 staining was restricted to bronchiolar epithelial cells and CGRP stained neuroendocrine cells in normal lungs (Figure 3g).

**Figure 3.**
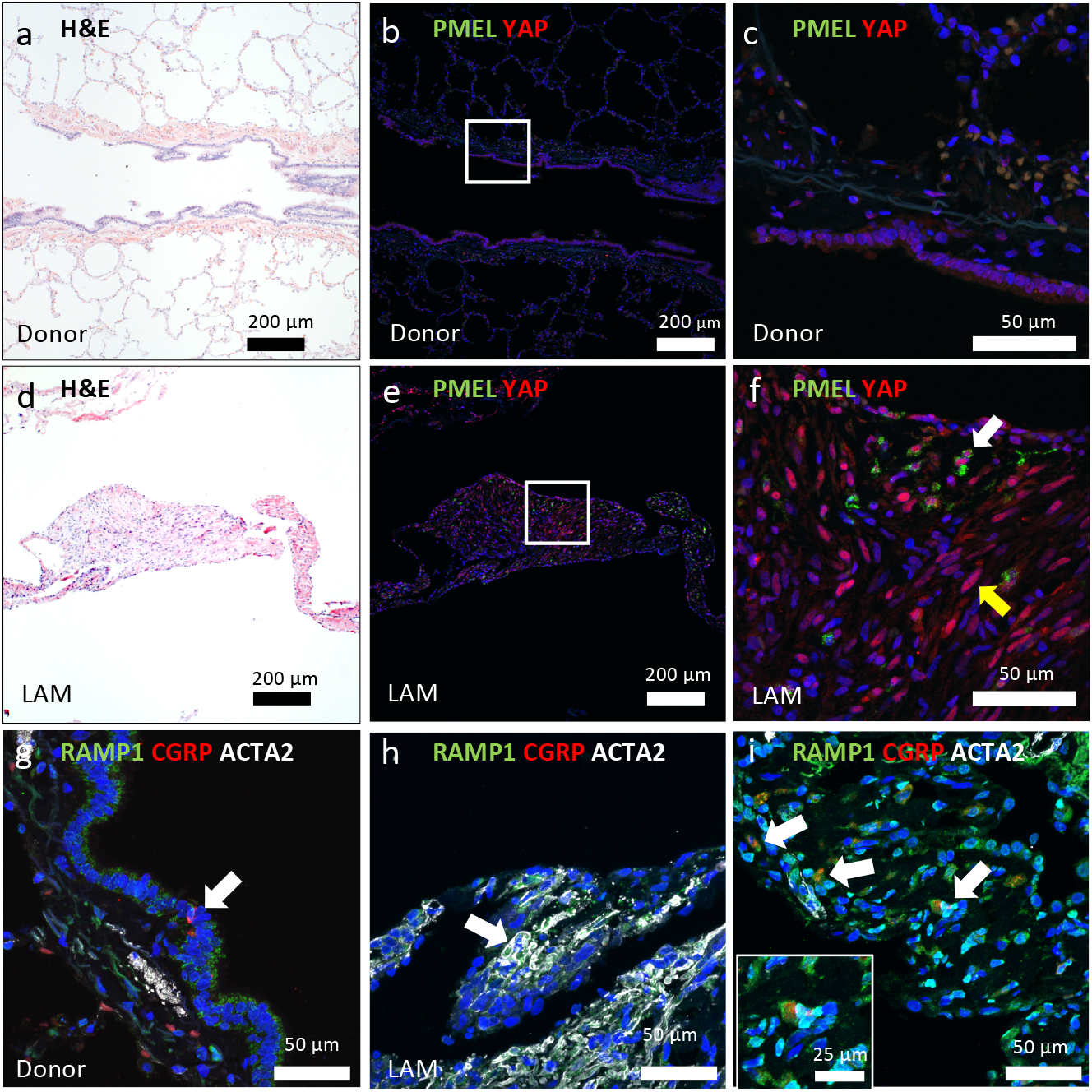
Detection of LAM^CORE^ signature genes and YAP. **(a)** Histology (H&E) of normal human lung bronchiole, surrounding smooth muscle and alveoli. **(b-c)** PMEL and YAP co-immunostaining showing weak nuclear staining of YAP in bronchiolar epithelium and absence in stroma in control lung, note non-specific reaction of red blood cells. **(d)** Histology (H&E) of LAM nodule and cyst. **(e-f)** PMEL and YAP immunostaining demonstrates nuclear YAP in PMEL positive cells (white arrow) and PMEL negative smooth muscle cells (yellow arrow). **(g)** RAMP1, CGRP, and ACTA2 co-immunostaining demonstrates RAMP1 expression in normal bronchiolar epithelium, RAMP1 and rare CGRP^+^ in neuroendocrine cells (white arrow) in control donor. **(h)** Abundant RAMP1 staining in ACTA2^+^ cells within LAM lesions. **(i)** RAMP1 and CGRP co-staining is seen in a subset of cells adjacent to ACTA2^+^ cells in a LAM lesion.

Since YAP and prostaglandin metabolic pathways have been reported to be upregulated in TSC and LAM (Li et al., 2017; Liang et al., 2014) and were predicted by scRNA-seq to be active in LAM^CORE^ cells (**Figure S4e)**, we assessed their expression in PMEL, ACTA2 positive LAM^CORE^ cells. Increased nuclear YAP was present in both PMEL^+^ cells and PMEL^-^ smooth muscle cells in LAM lung (Figure 3a-f). PTGER3 (Li et al., 2017) staining was detected in ACTA2-positive cells in LAM lesions (**Figure S10j**).

### The LAM^CORE^ secretome predicted by scRNA-seq

Since only a fraction (<20%) of the cells in LAM lesions contain mutations in TSC genes (Badri et al., 2013), it is clear that circulating and tissue resident lung cells are recruited and influenced by the LAM^CORE^ cells. To identify genes and processes active in LAM^CORE^ cells that influence adjacent pulmonary tissues, we identified LAM^CORE^ RNAs encoding secretome proteins using two databases, Human Protein Atlas (Thul et al., 2017) and VerSeDa (vertebrate secretome database) (Cortazar et al., 2017). A total of 162 LAM^CORE^ cell specific signature genes encode for secretome proteins (**Table S2**). Of these, 11 secretome proteins were exclusively found in LAM patient lungs and were not expressed in normal control lungs. IPA disease and functional enrichment analysis suggested that 9 of these 11 were associated with tumorigenesis in the female reproductive tract (*PMEL*, *IGSF21*, *C10orf90*, *NELL2*, *GOLM1*, *CPA6*, *PRLR*, *MMP11*, *CHI3L1*), and 6 of these 11 were associated with uterine carcinoma (*C10orf90*, *NELL2*, *GOLM1*, *CPA6*, *PRLR*, *MMP11*).

A number of known LAM serum biomarkers were identified in the LAM^CORE^ secretome including *VEGFD (FIGF), MMP2,* and *CTSK* (Ferri et al., 2004; Seyama et al., 2006). The expression of *PRLR* (prolactin receptor, also known as Secreted Prolactin Binding Protein), a newly predicted serum biomarker, is induced by ESR1 and PGR, and is capable of activating PI3K/AKT, ERK1/2 and TP53 signaling (Alkharusi et al., 2016; Terasaki et al., 2010). Genes encoding extracellular matrix associated proteins (i.e., collagen, glycosaminoglycans, proteoglycans, and endopeptidases) were the most enriched components in the LAM cell secretome and were associated with diverse LAM relevant functions including cell migration, angiogenesis, growth factor and estrogen responsiveness, and WNT, PI3K/AKT, P53 signaling pathways (Figure 4a).

**Figure 4.**
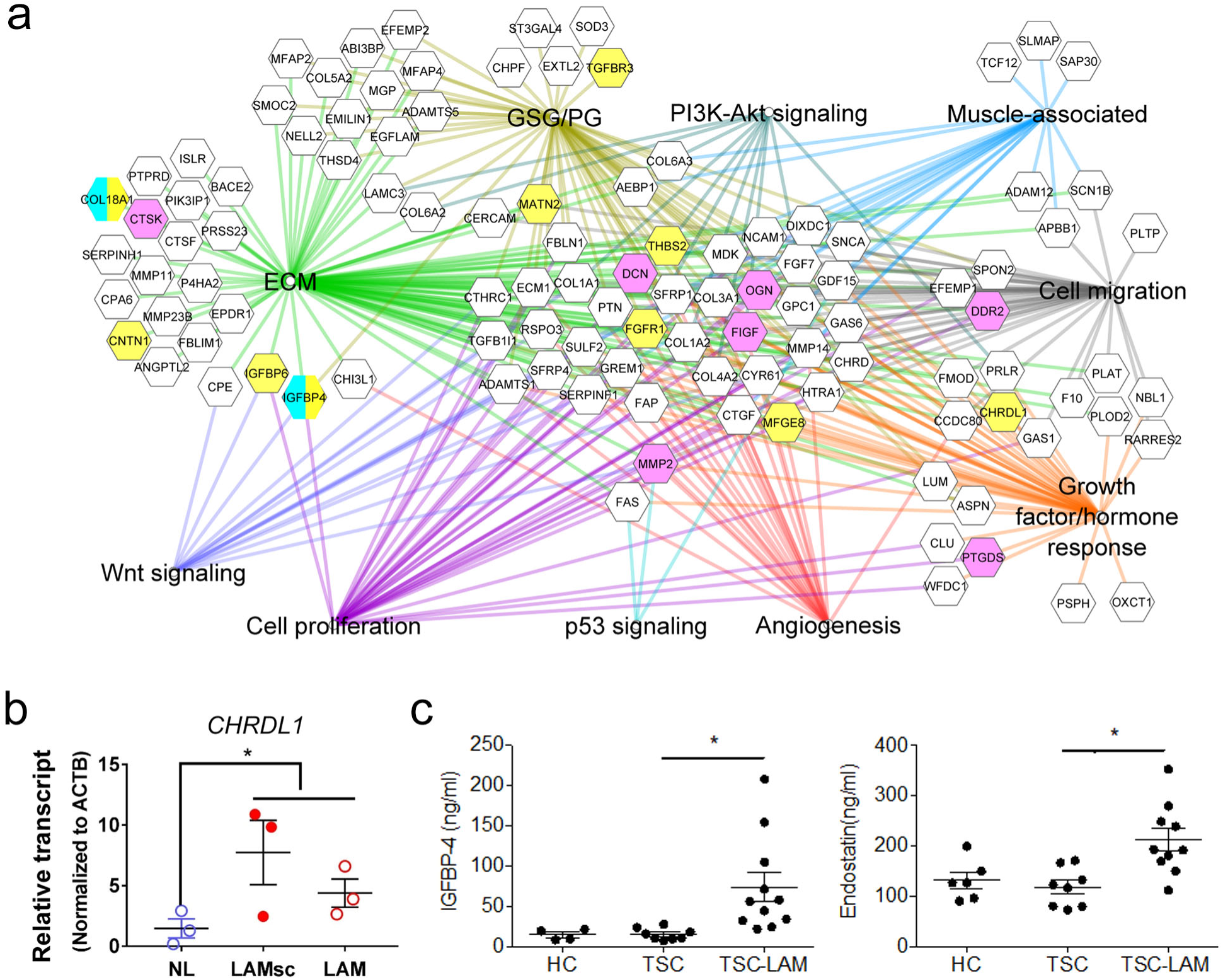
Prediction and validation of LAM^CORE^ cell “secretome”. **(a)** LAM^CORE^ signature genes encoding predicted secreted proteins and their enriched functional annotations are shown. Human Protein Atlas and VerSeDa databases were used to annotate secretome proteins. ToppGene suite was used to perform functional enrichment analysis. Network diagram was generated using Cytoscape (v3.7.1). Edge colors represent different bioprocess/pathways. Pink nodes are proteins reported in LAM literature. Proteins in yellow were validated by aptamer proteomics of LAM patient serum. Proteins in cyan were validated in LAM patient serum by ELISA. Proteins in yellow/cyan were validated by both aptamer proteomics and ELISA. ECM: extracellular matrix; GSG/PG: glycosaminoglycan/ proteoglycans. **(b)** *CHRDL1* RNA was measured in normal control (NL) and LAM tissues by RT-PCR. LAMsc: samples from LAM lungs used for scRNA-seq (LAM1, LAM3, and LAM4). LAM: samples from independently archived LAM lung tissues. Data are mean ± SEM. One-way ANOVA * P < 0.05. **(c)** ELISA analysis of serum levels of IGFBP-4 and Endostatin (COL18A1) in female TSC subjects with pulmonary LAM (TSC-LAM), TSC women without pulmonary LAM (TSC), and healthy subjects (HC). *, P < 0.05; one-way ANOVA.

RNA analysis, immunofluorescence staining of tissues, serum aptamer proteomics (Gold et al., 2010) and ELISA assays were used to validate the predicted LAM^CORE^ secretome. Aptamer proteomics of serum from female TSC-LAM patients identified increased levels of proteins encoded by LAM^CORE^ signature genes compared to women with TSC without LAM, including CHRDL1, CNTN1, COL18A1, FGFR1, IGFBP4 (Valencia et al., 2001), IGFBP6, MATN2, MFGE8, TGFBR3, and THBS2 (Figure 4a). Increased expression of *CHRDL1* in LAM lesions was validated via qRT-PCR (Figure 4b). Predicted novel LAM secreted proteins, IGFBP-4 and endostatin (derived from proteolysis of COL18A1) were increased in serum from female TSC-LAM patients compared TSC without LAM, or healthy women by ELISA (Figure 4c).

### LAM^CORE^ cell origins predicted by scRNA-seq analysis

We compared LAM scRNA-seq data with RNA profiles from multiple organ signatures in the HumanGeneAtlas (Su et al., 2004), Bodymap (Hishiki et al., 2000) and Molecular Signatures Database (MSigDB) (Liberzon et al., 2015) to predict the tissue of origin for LAM^CORE^ cells. Genome-wide, transcriptomic similarities were assessed using unbiased hierarchical clustering. Pseudo-bulk RNA profiles derived from scRNA-seq data from LAM^CORE^, control lung and AML (angiomyolipoma *ACTA2*^+^ cell cluster) were compared. Pseudo-bulk LAM^CORE^ RNA profiles co-clustered with *ACTA2*^+^ cells from AML (AML.*ACTA2*^+^) and were closest to normal uterus, ovary, adipocytes and neural ganglia (Figure 5a). As expected, pseudo-bulk data from control lung clustered most closely with normal lung (Figure 5a; **Figure S12**). Functional enrichment analysis of the LAM^CORE^ signature genes (n=777) using ToppGene suite showed they were most closely associated with uterine myometrium (**Figure S13**; **Table S6**).

**Figure 5.**
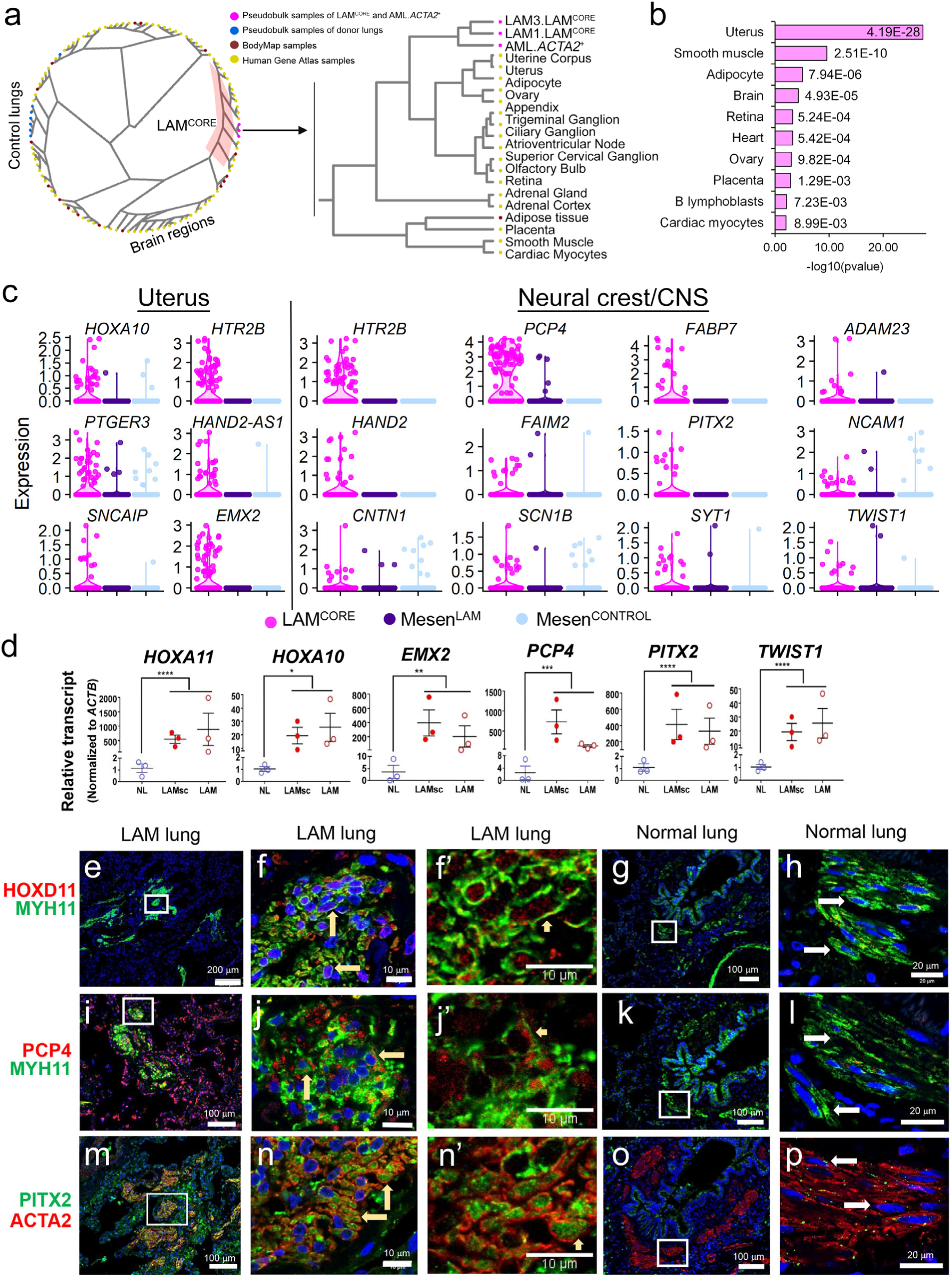
Transcriptomic similarity of LAM^CORE^ cells to other cells and organs. **(a)** Hierarchical clustering analysis of pseudo-bulk profiles (n=10) of LAM^CORE^, AML.*ACTA2*^+^, and normal lung cells, microarray RNA profiles (n=84), and bulk RNA-seq data (n=16) demonstrated co-clustering of LAM^CORE^ transcriptomic profiles with RNA samples from uterus, smooth muscle, and adipocyte. AML.*ACTA2*^+^: mesenchymal cells from AML patient. Left panel shows radial presentation of the hierarchical tree; right panel shows the LAM^CORE^ subtree. **(b)** Human tissues whose signatures were most associated with the LAM^CORE^ signature. Significance of association was tested using Fisher’s exact test. RNA profiles of human tissues were from HumanGeneAtlas (HGA) dataset. **(c)** Violin plots depict selective expression of uterine and neural crest/CNS genes in LAM^CORE^. Mesen^LAM^: LAM-associated mesenchymal cells; Mesen^CONTROL^ : mesenchymal cells from normal lung. CNS: central nervous system. **(d)** LAM^CORE^ marker genes were measured in normal control (NL) and LAM tissues by RT-PCR. LAMsc: LAM lungs used for scRNA-seq. LAM: independently archived LAM lungs. Data are mean ± SEM. One-way ANOVA. *, P < 0.05; **, P< 0.01, ***, P < 0.005; ****, P < 0.001. **(e-p)** Lung tissues from LAM patients and control donors were immunostained and imaged by confocal immunofluorescence microscopy. Nuclear HOXD11 and PCP4 was detected in MYH11^+^ cells in LAM lung lesions in LAM lung and was absent in control lung (e-l). PITX2 was present in ACTA2^+^ cells in LAM lesions in LAM lung and was absent in control lung (m-p). Panels f’, j’, and n’ show staining without DAPI. Yellow arrows indicate positive staining, and white arrows indicate non-positive staining.

Considering that genome-wide correlations and functional enrichments are highly influenced by the most abundantly expressed genes from each tissue, we assessed transcriptomic similarities using tissue specific signature genes. Tissue selective signature genes were identified for 84 different human tissues (**Figure S13b**) and compared with the LAM^CORE^ signature (Figure 5b; **Tables S7-8**). Uterus, smooth muscle, adipocyte, brain regions and retina were identified as tissues with the most similarity to the LAM^CORE^ signature via Fisher’s exact test (Figure 5b; **Table S7**). Representative uterine and neural crest signatures were significantly enriched in LAM^CORE^ but not in LAM associated mesenchymal cells or donor mesenchymal cells (Figure 5c).

RNA analysis and immunofluorescence staining were used to validate a number of newly identified LAM^CORE^ cell signature genes that are typically selectively expressed in uterus and neural crest and not in lung. Uterine (*HOXA11, HOXD11, EMX2, RAMP1, PTGER3, PGR*) and brain/neural crest (*PCP4, UNC5D, KCNAB1, PITX2 and TWIST1*) associated genes were abundantly expressed in the pulmonary LAM lesions but barely detectable in normal female lungs (Figure 5d; **Figure S10**). Nuclear HOXD11 (Homeobox D11) and PITX2 (Paired Like Homeodomain 2) were present in ACTA2^+^ cells in LAM lung lesions and absent from normal female lung (Figure 5e-h**, 5m-p**). Similarily, PCP4 (Purkinje Cell Protein 4) was detected in cytoplasm of MYH11^+^ cells in LAM lung lesions and was absent in control lung (Figure 5i-l). RAMP1 (normally expressed in endometrium and smooth muscle) and ACTA2 were co-expressed in LAM nodules but not in control lung (Figure 3g-i).

### Identification of LAM^CORE^ cells in uterus from a patient with LAM

Uterine LAM lesions have been reported in rare patients with pulmonary LAM (Hayashi et al., 2011). To determine whether uterine LAM cells share the unique LAM^CORE^ cell transcriptome identified by pulmonary LAM single-cell and single-nuclei RNA-seq, we performed snRNA-seq on uterine tissue directly obtained from the operating room at the time of hysterectomy for a sporadic LAM patient and a normal female donor. Gross inspection and histopathology revealed multiple well circumscribed, firm nodules with a whorled trabecular pattern bulging from the myometrial surface, consistent with leiomyomas. No macroscopic LAM lesions were identified, consistent with previous studies reporting that most uterine LAM lesions are microscopic and localized in the subserosal myometrium (Hayashi et al., 2011). Based on these observations, uterine tissue containing subserosal myometrium was selected for snRNA-seq in an attempt to enrich for LAM cells. Unbiased clustering analysis identified a distinct cluster of cells sharing RNA expression patterns with pulmonary LAM^CORE^ cells, including known markers *PMEL* and *FIGF*, and novel LAM^CORE^ markers *UNC5D*, *KCNC2*, *NECAB2*, and *KCNAB1* (Figure 6a; **Figure S15**). In addition, uterine myocyte, epithelial, immune, endothelial and several stromal cell subtypes were identified (Figure 6a; **Figure S15**). Endometrial epithelia (expressing high level of *KRT8, KRT18, EPCAM, CLDN3*) and stromal cells (expressing high level of *MME, FN1, COL3A1, HOXA10, VIM*) were defined based on the gene markers from the human and mouse uterus altas (Mucenski et al., 2019; Wu et al., 2018). Hierarchical clustering of major cell types from LAM lung and LAM uterus demonstrated close correlation of LAM^CORE^ cells from both tissues (Figure 6b).

**Figure 6.**
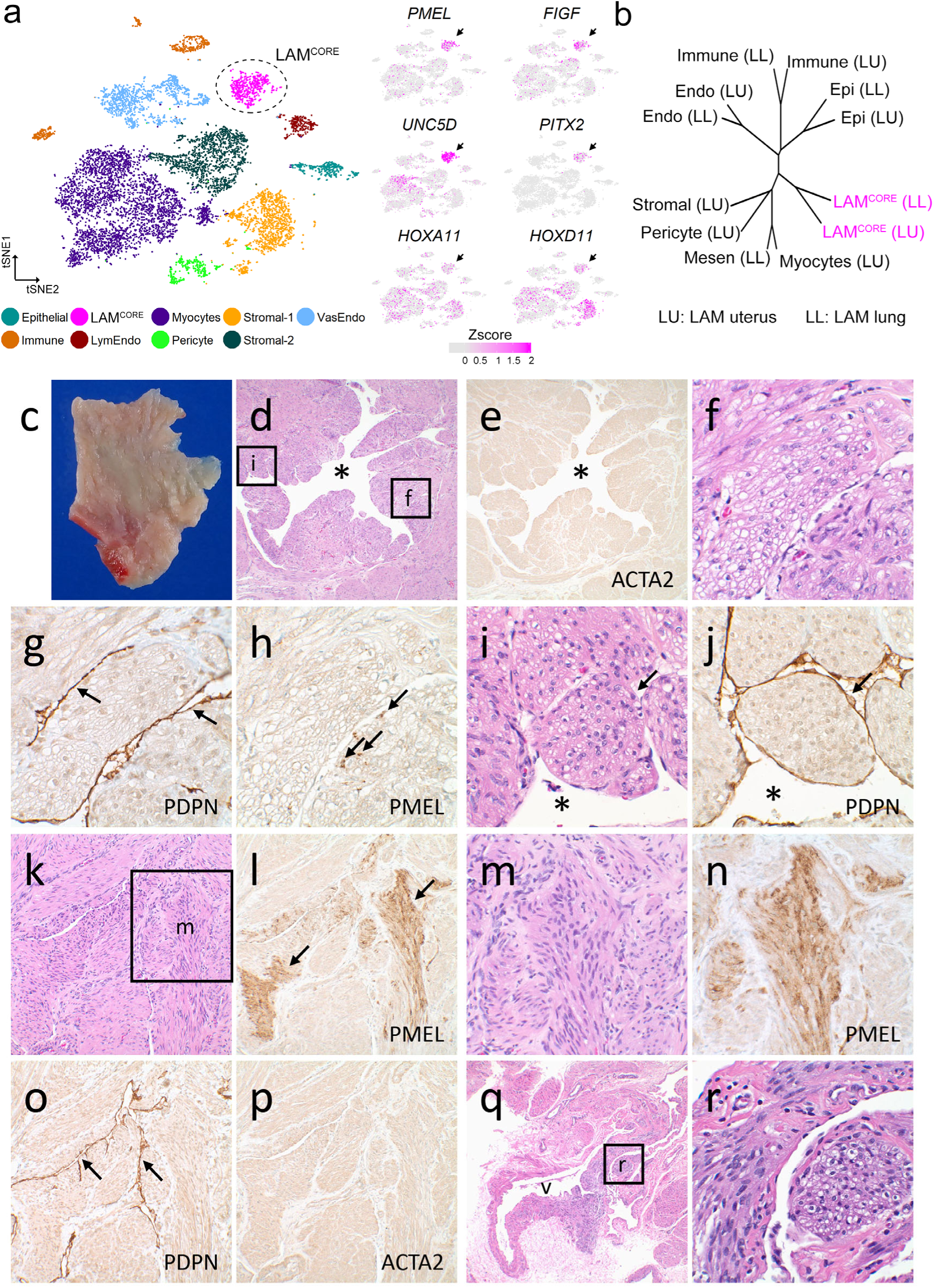
Expression of LAM^CORE^ signature genes in LAM uterus. **(a)** snRNA-seq analysis of LAM uterus identified a cluster of cells expressing LAM^CORE^ signature genes. Left panel shows cells in tSNE plot; dashed line encircles uterine LAM^CORE^ cells (magenta). Right panels show expression of LAM^CORE^ signature genes in tSNE plots of LAM uterus; black arrows indicate uterine LAM^CORE^ cell cluster. **(b)** Hierarchical clustering of major cell types of single cells/nuclei from LAM lung and LAM uterus. Epi: epithelial cells; Endo: endothelial cells; Mesen: mesenchymal cells; Immune: immune cells. LL: scRNA-seq data from LAM lungs; LU: snRNA-seq data from LAM uterus. Text in magenta labels LAM^CORE^ cells. **(c-r)** LAM lesions in uterus of LAM patient. (c) Fresh uterine tissue from LAM patient showing subserosal myometrium adjacent to tissue used for snRNA-seq analysis. No macroscopic LAM lesions were identified. (d-j) LAM lesions in the uterus identified microscopically resembled lung LAM lesions. Uterine LAM lesions were comprised of variably dilated spaces (d-e*) surrounded by bundles and nodules of smooth muscle actin (ACTA2) stained (e) spindled and epithelioid LAM cells with eosinophilic to clear cytoplasm (f). Lesions were lined by podoplanin (PDPN) stained lymphatic endothelial cells (g, arrows), with a variable subpopulation of LAM cells staining with HMB-45 antibody for PMEL (h, arrows). LAM cell clusters (LCC) (i, arrow) surrounded by a monolayer of PDPN stained lymphatic endothelial cells (j, arrow) within lymphatic spaces (i-j*). (k-r) A second uterine LAM lesion (k) with abundant PMEL immunopositive cells (l, arrows). Higher power images demonstrate bundles of spindled LAM cells (m) with characteristic granular cytoplasmic PMEL staining (n). Compressed lesions were lined by PDPN stained endothelial cells (o, arrows), around the bundles of LAM cells that were diffusively stained for ACTA2 (p). Vascular involvement by LAM cells was present with infiltration of the vessel (q, labeled v) wall by spindled and epithelioid LAM cells (r). Original magnification: 100x (d-e, q); 200x (k-l, o-p); 400x (m-n); 600x (f-j, r).

Uterine and neural crest associated genes identified in pulmonary LAM^CORE^ cells were compared using scRNA-seq data from the human LAM uterus and normal human/mouse uterus (Mucenski et al., 2019). Most neural crest and uterine associated genes selectively expressed in pulmonary LAM^CORE^ cells were expressed in the LAM cell cluster from the human LAM uterus (Figure 6a; **Figure S15**). A detailed comparison of common vs unique signature genes in pulmonary vs uterine LAM^CORE^ cells is shown in **Figure S15**. The conserved markers are largely associated with neural crest cell migration and neuronal differentiation (*PITX2*, *TWIST1*, *HAND2*, *UNC5D*, *DCX*). Compared with uterine LAM^CORE^ cells, pulmonary LAM^CORE^ cells were more closely associated with extracellular matrix components, cell movement and muscle elements (*MMPs*, collagens, laminins, ect); while uterine LAM^CORE^ cells expressed more transmembrane transport and brain/neural crest genes, including multiple solute carrier family members, potassium voltage-gated channels, cholinergic receptors, and glutamate ionotropic receptors; many are associated with neurodegenerative disorders (**Figure S15**). Importantly, myocytes and stromal cells in the normal human and mouse uterus (**Figure S14**) expressed both neural crest and uterus associated genes. Immunofluorescence staining of uterine marker HOXD11 and neural crest marker PCP4 demonstrated their expression in normal human uterus (**Figure S11**).

The presence of LAM^CORE^ cells within the uterine LAM tissues used for snRNA-seq was demonstrated by microscopic and immunohistochemical analysis of contiguous histologic sections (Figure 6c-r). Multifocal uterine LAM lesions were morphologically and phenotypically similar to pulmonary LAM lesions in many ways: uterine LAM lesions were comprised of spindled and epithelioid LAM cell bundles and nodules surrounding variably dilated lymphatic channels lined by podoplanin (PDPN) stained lymphatic endothelial cells (Figure 6c-r); LAM cell clusters composed of LAM cells surrounded by an abundance of lymphatic endothelial cells were observed in uterine LAM lesion (Kumasaka et al., 2005) (Figure 6i-j); uterine LAM lesions were stained diffusely for ACTA2 and were focally positive for PMEL (Figure 6e-p). Together, these data demonstrate that LAM cells in the lung and uterus are morphologically indistinguishable and share similar gene expression profiles. In summary, both bioinformatic analyses and experimental evidence strongly support uterus as the likely origin of LAM^CORE^ cells.

## DISCUSSION

Recurrence of pulmonary LAM following lung transplantation led to the realization that LAM is a low-grade metastatic neoplasm rather than an interstitial lung disease arising from expansion of tissue resident mesenchymal cells (Carsillo et al., 2000; McCormack et al., 2012). An understanding of the source of cells that infiltrate the lung has remained elusive, however, as have the mechanisms of tumorigenesis, metastasis, invasion, and cystic remodeling (Henske and McCormack, 2012; Krymskaya and McCormack, 2017; McCormack et al., 2012). In the present study, scRNA-seq analysis of LAM lesions identified unique, non-pulmonary mesenchymal cells, which we termed LAM^CORE^ cells, that share a high degree of transcriptomic similarity to the LAM^CORE^ cells in uterus of an S-LAM patient and with myometrial and stromal cells (express high level of *MME, FN1, COL3A1, HOXA10*) from normal human and mouse uteri (Mucenski et al., 2019; Wu et al., 2018). A prominent neural crest signature was also identified in pulmonary and uterine LAM^CORE^ cells and in normal uterus. scRNA-seq profiling, together with aptamer proteomics of patient serum, qPCR, immunofluorescence confocal microscopy and serum ELISA identified novel secreted proteins, cell specific signature genes, and altered bioprocesses and signaling pathways characteristic of LAM^CORE^ cells.

LAM^CORE^ cells express unique gene signatures that are driven by the cell autonomous effects of constitutive mTOR activation and by interactions with neighboring and recruited cells in their pulmonary niche. The transcriptomic profile of the LAM^CORE^ cells is not consistent with that of any known pulmonary cell, either in the literature or the current study, but share closest transcriptomic similarity with normal uterine myocytes and neural crest. The congruity of gene expression in uterine and pulmonary LAM^CORE^ cells is consistent with a uterine source for pulmonary LAM cells, and is supported by the remarkable penetrance of myometrial tumors (100%) with spontaneous pulmonary metastasis (50%) in the mouse model of myometrium specific *Tsc2* deletion (Prizant et al., 2013; Prizant et al., 2016). The strong neural crest transcriptomic signature in both mouse and human suggests that uterine LAM cells may themselves have had developmental origins in the neural crest. Together with the clinical features of female gender restriction, pelvic predominant distribution of lymph node involvement, hormonal responsiveness, and prior case reports of uterine LAM provide suggestive evidence that LAM is a low grade metastatic uterine tumor. The occurrence of LAM in men indicates that other sources for pulmonary LAM cells must also exist, perhaps from other sites populated by neural crest cells bearing TSC mutations. The occurrence of PEComas in other locations within the male and female genitourinary tract (e.g., kidney, bladder, ovary, prostate, etc.), gastrointestinal tract (e.g., stomach, small intestine, colon, liver etc.), soft tissue and bone suggest additional potential candidates for the primary tumor in both men and women with LAM (Martignoni et al., 2008).

It seems rather implausible that such an obvious source for LAM as the uterus could have been largely overlooked for the 80 and 100 years since S-LAM and TSC-LAM (respectively) first appeared in the literature. Indeed, there have only been a few dozen cases of uterine LAM reported individually and in one small series (Hayashi et al., 2011). It is important to note that LAM cells often blend seamlessly into the myometrial tissue of the uterus, and can easily be missed by pathologic examination. In addition, uterine LAM lesions are subserosal and sparse in S-LAM specimens (compared to diffuse and abundant in TSC-LAM uteri), often requiring step-sectioning for identification (Hayashi et al., 2011). For these reasons, we believe identification of LAM in the uterus has likely been underreported, and that use of markers such as HMB-45 and podoplanin should be considered in the pathologic examination of LAM uterine specimens, even in the absence of convincing morphological features of LAM. A number of new, unique LAM cell markers identified herein by scRNA-seq, such as HOXD11, PITX2, may serve as useful diagnostic tools in the future.

A model for LAM has been proposed in which somatic mutations in TSC2 lead to mTOR activation, cellular proliferation and VEGF-D-dependent lymphangiogenic programs that drive recruitment of lymphatic endothelial cells, lymphatogenous spread to the lung, and cystic lung remodeling (Henske and McCormack, 2012; Kumasaka et al., 2004; Kumasaka et al., 2005). Evidence for this concept has been limited to histological and immunohistochemical studies that demonstrate LAM cell clusters enveloped by a monolayer of lymphatic endothelial cells within the lumen of the thoracic duct, conducting lymphatics and chylous effusions, and within chaotic lymphatic channels lined by VEGFR-3 (FLT4) expressing lymphatic endothelial cells in primary lung and uterine lesions of LAM patients (Kumasaka et al., 2004; Kumasaka et al., 2005). The findings that both lung and uterine LAM^CORE^ cells show predominant *VEGF-D* expression, that *FLT4* (VEGFR-3) RNA is highly expressed in lymphatic endothelial cells, and that *VEGFC* is mostly expressed in vascular endothelial cells, support an important role for lymphangiogenesis in the pathogenesis of LAM. Two of the predicted LAM^CORE^ signature genes are thought to play key roles in lymphatic specification (Iwasaki et al., 2019), NR2F2, a nuclear receptor that was recently identified by a genome-wide association study (GWAS) as a modifying genetic locus in LAM (Kim et al., 2019), and EP3 (PTGER3), the receptor for prostaglandin E2, were among the LAM signature genes identified that could well be driving lymphangiogenic programs (Li et al., 2017) (**Table S2**).

scRNA-seq of LAM tissues demonstrated a robust activation of mTOR pathway in LAM^CORE^ cells, consistent with the role of TSC gene mutations. Present analysis identified other signaling pathways that may influence cellular proliferation, migration, invasion and transdifferentiation of LAM^CORE^ cells as shown in **Figure S4**. LAM^CORE^ cells express multiple growth factors, including *VEGFD, PDFGRA/B, FGF7, IGFBP2/4/6,* and *TGFB1I1* that likely mediate active cross-talk between pulmonary LAM^CORE^ cells and surrounding cells (Figure 4a; **Table S2**). RNAs encoding the LAM^CORE^ cell secretome provide insights into potential mechanisms by which LAM cells orchestrate tissue destruction and shape the cellular microenvironment in the LAM lung. Genes encoding extracellular matrix (ECM), including glycosaminoglycans, proteoglycans, diverse collagens, proteolytic enzymes (MMPs and ADAMs), and a number of matricellular proteins including thrombospondins, fibulins, SPARC, CYR61 and CTGF, were the most enriched functional classes of secretome proteins in LAM^CORE^ cells (Nikitovic et al., 2014). The finding that LAM^CORE^ cells are predominant source of VEGF-D (FIGF) expression in the lung validates its use as a biomarker of LAM cell viability and mTOR pathway activation (Young et al., 2013). Multiple other serum biomarker candidates were identified from the LAM^CORE^ cell secretome analysis (Figure 4a; **Table S2**). We validated the increased expression of Col18A1 (endostatin) and IGFBP-4 in serum of patients with TSC-LAM. Endostatin, an anti-angiogenic cleavage product of matrix protein Col18A1 (Wickstrom et al., 2005) may bias vasculogenesis toward lymphatic lineages, and IGFBP-4, a protein that binds and enhances the action of IGF-1 and IGF-2 (Valencia et al., 2001) may provide growth advantages to LAM cells.

LAM^CORE^ cells from LAM lungs expressed a unique panel of signature genes expressed in uterine LAM cells and in normal human and mouse uterus but not in normal human lung. LAM^CORE^ cells and normal myometrial cells shared transcriptomic similarity which are regulated by female sex hormones, including estrogen, progesterone and prolactin receptors (Prizant et al., 2016; Terasaki et al., 2010). scRNA-seq analysis of normal human and mouse uterine cells demonstrated a close relationship of LAM^CORE^ signatures with uterine myocytes. Brain/neural crest RNAs identified in pulmonary LAM^CORE^ cells were also enriched in myocytes from normal human and mouse uteri and in LAM^CORE^ cells in the uterus from a LAM patient. Since neural crest cells are capable of differentiating into a variety of mesenchymal cell types, including smooth muscle cells in several organs (Delaney et al., 2014; Wang et al., 2012), uterine neural crest is a likely source of the LAM^CORE^ cells.

There are several limitations in the present study, most related to the rarity of LAM disease and limited availability of fresh tissue samples. The yield of mesenchymal cells (including LAM cells) isolated from the tissues for scRNA-seq analysis of all lung specimens was relatively low, perhaps related to incomplete liberation of the cells from matrix or effects of the digestion protocol on viability.

In summary, we used scRNA and snRNA sequencing to identify a unique population of LAM^CORE^ cells in both LAM lung and uterus. Bioprocesses and signaling pathways which were active in pulmonary LAM^CORE^ cells were most similar to those in uterus and were distinct from those in normal lung, supporting the concept of the uterus as the source of LAM^CORE^ cells. The present findings provide new insights into the characteristics of LAM cells and predict signaling networks by which LAM^CORE^ cells infiltrate, recruit and interact with other cells in the lungs of LAM patients. Single cell secretome analyses identified a number of novel LAM cell-specific secretome proteins that hold promise as potential biomarkers for diagnosis or prognosis and therapeutic targets. More immediate products of this work could include the rapid development of new immunohistochemical approaches for the diagnosis of LAM based on novel markers (such as PITX2 or HOXD11), refinement of the approach to pathological analysis of uterine LAM lesions and development of new serum biomarkers. Present data support further study of the utility of hysterectomy for source control in some patients with early LAM or at risk for LAM, such as young women with TSC not considering future pregnancy, and redoubled efforts to design and execute hormonal therapy trials for premenopausal patients with LAM. Finally, advancing the understanding of LAM pathogenesis as a unique, monogenic model of malignancy(Krymskaya and McCormack, 2017) may yield broader insights into the biology of cancer.

## METHODS

### Human samples

Lung tissues from LAM patients were obtained from the National Disease Research Interchange (NDRI), with the assistance of The LAM Foundation. Female donor lung tissues were obtained from LifeCenter Organ Donor Network-Cincinnati. Uterine tissue from a ‘normal’ donor was obtained from the Univeristy of Cincinnati, and the LAM uterine sample was obtained from St Vincent Women’s Hospital, Indianapolis, Indiana. The Institutional Review Board of the University of Cincinnati College of Medicine approved all human-relevant studies. Single-cell RNA-seq (scRNA-seq) was performed using the 10x Chromium platform on three lung explants from LAM patients undergoing lung transplantation (LAM1, LAM2, and LAM3, **Table S1**). Single nuclei RNA-seq (snRNA-seq) was performed on a fourth LAM explant (LAM4), in an attempt to improve the representation of mesenchymal cells in the analysis. LAM2 tissue was obtained from a patient taking Sirolimus while donors of LAM1, LAM3 and LAM4 were not being treated with Sirolimus at the time of transplant. We also performed scRNA-seq on a renal AML resected from a patient with a sporadic AML (S-AML), and snRNA-seq on uterine tissue obtained at hysterectomy from an S-LAM patient. As controls, we conducted scRNA-seq on the explanted lung of a brain dead, beating-heart, organ donor (female, age 31) and snRNA-seq on “normal uterine tissue” from a uterus resected for cervical cancer (age 29), and obtained scRNA-seq data of six additional female donor lungs from Gene Expression Omnibus (GEO) (Reyfman et al., 2018). Additional archival LAM lung specimens from National Disease Research Interchange (NDRI) were used for IHC and RT-PCR in cross validation experiments.

### TSC1/TSC2 mutation analysis

High read depth (500-1,000x) massively parallel sequencing was performed to identify TSC1/TSC2 mutations in LAM and angiomyolipoma samples, as described previously (Giannikou et al., 2019). Briefly, 100 ng DNA was subjected to hybrid capture (Agilent SureSelect platform, Santa Clara, CA, USA) for the entire genomic extent of TSC1 and TSC2, the resulting library was sequenced (Illumina platform, San Diego, CA, USA), followed by custom computational analysis for sequence variants. Candidate TSC1/TSC2 variants were reviewed in IGV, and were validated by PCR-based amplicon massively parallel sequencing.

### Single cell isolation from LAM lung and AML kidney

The enzyme mix for tissue digestion was made up in DPBS (Thermo Fisher #14190) containing 6 mg/mL collagenase A (Sigma #10103578001), 4.3 U/mL elastase (Worthington #LS002292), 9 u/mL dispase (Worthington #LS02100), 100 µg/mL soybean trypsin inhibitor (Roche #10109886001), 125 U/mL DNAse (Applichem #A3778) and 5 mM CaCl2. DPBS with 0.04% BSA (Sigma #A8806) was used for washing and re-suspending cells. An area of the tissue predicted to be enriched for LAM cells was cut into small pieces, transported in ice-cold DPBS and minced thoroughly in a petri dish on ice for 5 min. Tissue was divided into 13 mg portions and placed into tubes with 1 mL enzyme mix and incubated on ice for 40 min with shaking every 2 min and trituration (10 times) every 3 min. After 40 min of digestion, tissue was allowed to settle on ice for 1 min. The supernatant containing liberted cells was carefully removed and applied to a 30 µM MACS filter (Miltenyi #130-098-458) in a 15 mL conical tube, followed by rinsing with 12 mL ice-cold PBS/BSA. The flow-through was divided into two 15 mL conical tubes, and each volume brought to 14 mL with ice-cold PBS/BSA. The procedure was repeated for the remaining tissue pieces, and the cells obtained were combined with the first run. The viability of the cell preparation was assessed with trypan blue, and the cell concentration was adjusted to 1,000 cells/µL for single cell analysis with 10x chromium.

### Single nuclei isolation from LAM lung and uterus

LAM lung tissue was snap frozen in liquid nitrogen and stored at −80 °C prior to isolation. Tissue (50 mg) was minced thoroughly on ice for 5 min using razor blades and forceps and placed in 2 mL ice-cold EZ nuclei lysis buffer (Sigma #NUC101-1KT) in a Dounce homogenizer (Sigma #D8938). For uterine tissue, nuclei were isolated as described by Humphrey *et al*. (DOI: dx.doi.org/10.17504/protocols.io.nahdab6). Tissue pieces were dounced 40 times with loose pestle followed by 20 times with a tight pestle and transferred to a 15 mL conical tube to which 2 mL of Nuclei EZ lysis buffer was added. Nuclei were incubated for 10 min on ice, with trituration (10 times) every 2 min and pelleted by centrifugation at 500 g at 4 °C. The supernatant was discarded and nuclei were re-suspended in 4 mL ice-cold Nuclei EZ lysis buffer and incubated for 10 min on ice with trituration. Nuclei were centrifuged at 500 g for 5 min at 4 °C, the supernatant was discarded, and nuclei were re-suspended in 4 mL ice-cold NSB containing 1% BSA/DPBS and 0.2 U/mL RNAse inhibitor (Takara #2313A) and passed through a 30 µM filter. Nuclei were pelleted at 500 g for 5 min at 4 °C and the supernatant discarded. Nuclei were suspended in 600 µL ice-cold NSB. An aliquot (50 µL) was taken as a negative control for FACS. To the remining aliquot, 3 µL of Hoescht stain (Invitrogen, H3570) was added, mixed by trituration and shaken for 5 min on ice. FACSAria, with a 70 µM nozzle size, was used to sort DAPI+ stained nuclei. Nuclei were collected in an Eppendorf tube containing 500 µL of NSB and pelleted at 2500 RPM for 5 min at 4 °C. Supernatant was removed and the nuclei re-suspended in 50 µL ice-cold NSB. Nuclei were quantified using a hemocytometer with trypan blue and the concentration was adjusted to 1000 nuclei/µL for 10x Chromium.

### Pre-processing of single-cell/single-nuclei RNA sequencing data

Read alignment and gene-expression quantification of human scRNA-seq and snRNA-seq data were performed using CellRanger pipeline (version 2.0.0 and version 3.0.2, 10x Genomics) (Zheng et al., 2017). For scRNA-seq alignment and quantification, CellRanger pre-built human (hg19) reference package was used. For snRNA-seq alignment and quantification, a CellRanger compatible human (hg19) “pre-mRNA” reference package was generated according to Cell Ranger’s instructions (https://support.10xgenomics.com/single-cell-gene-expression/software/pipelines/latest/advanced/references). scRNA-seq data (Reyfman et al., 2018) of six female human lungs were downloaded from GEO (GSE122960). For all scRNA-seq data, cells with at least 500 expressed (UMI>0) genes and less than 10% of UMIs mapping to mitochondrial genes were included for downstream analysis. For snRNA-seq data of LAM4, cells with at least 300 expressed (UMI>0) genes and less than 10% of UMIs mapping to mitochondrial genes were included. For snRNA-seq data of LAM uterus and normal uterus, cells with at least 1,000 expressed (UMI>0) genes and less than 10% of UMIs mapping to mitochondrial genes were included. For scRNA-seq and snRNA-seq data, genes expressed in at least two cells in an individual dataset were included. Downstream analysis was performed in R 3.5.0 using custom scripts, SINCERA (Guo et al., 2015; Guo and Xu, 2018), and Seurat (Butler et al., 2018; Satija et al., 2015) (versions 2.3.4 and 3.0.0).

### scRNA-seq/snRNA-seq data analysis

The following pipeline was setup to analyze scRNA-seq/snRNA-seq data from an individual sample. The expression of a gene in a cell was measured by its UMI counts in the cell normalized by the total number of UMIs in the cell, multiplied by 10,000, added by 1, and then taking the natural log. The FindVariableGenes function in Seurat2 (Butler et al., 2018) was used to identify highly variable genes. The expression of highly variable genes was used for dimension reduction using principal component analysis. Cell clusters were identified using Louvain-Jaccard algorithm (Shekhar et al., 2016) using the principal components with largest variances. Cluster specific differentially expressed genes were identified using a binomial based (Shekhar et al., 2016) and a negative binomial based differential expression tests implemented in the SINCERA pipeline (Guo et al., 2015; Guo and Xu, 2018; Yu et al., 2013). The negative binomial test based method is based on sSeq (Yu et al., 2013) and uses the total UMI counts for each cell divided by the median UMI counts per cell as relative library size as suggested in CellRanger (Zheng et al., 2017). Genes with false discovery rate of *p* value <0.1 and fold change >=2 in both tests were considered significant. For the binomial test based method, effect size (Shekhar et al., 2016) was considered as fold change. Clusters were assigned to cell types based on the expression of known marker genes and functional enrichment analyses using ToppGene suite (Chen et al., 2009) with cluster specific differentially expressed genes as input.

### Integrative analysis of scRNA-Seq of LAM samples

Pre-processed scRNA-seq data from the two LAM samples (LAM1 and LAM3) were aligned using the canonical correlation analysis in Seurat 2 (Butler et al., 2018). Cell clusters were identified using Louvain-Jaccard algorithm (Shekhar et al., 2016). Clusters were assigned to cell types (endothelial, epithelial, immune, and mesenchymal cell subtypes, and LAM^CORE^) based on the expression of marker genes and functional enrichment analyses using ToppGene suite (Chen et al., 2009) with cluster specific differentially expressed genes as input.

The LAM^CORE^ signature genes were defined on the basis of one of two criteria: 1) genes differentially expressed in LAM^CORE^ cluster (DE), i.e. genes that passed both binomial and negative binomial based tests at FDR (Storey and Tibshirani, 2003) (false discovery rate) <0.1, fold change >=2, and expression frequency >=10% in the LAM^core^ cluster (DE genes, n=760); or 2) genes uniquely expressed in LAM cluster (UE), i.e., genes with expression sensitivity >=80% and expression frequency >=10% in the LAM^CORE^ cluster in either scRNA-seq sample (UE genes, n=36).

### Integrative analysis of LAM scRNA-seq and snRNA-seq data

We first analyzed the pre-processed LAM sc and snRNA-seq data seperately, identified cell clusters using the Louvain-Jaccard method (Shekhar et al., 2016), detected cluster specific differentially expressed genes using the binomial and negative binomial tests based methods, and assigned clusters to major cell types based on the expression of marker genes and functional enrichment analysis using ToppGene suite (Chen et al., 2009). We then aligned the separately analyzed LAM scRNA-seq and snRNA-seq data using Harmony (Korsunsky et al., 2018) with expression values imputed using ALRA (Linderman et al., 2018). Cell clusters were identified using Louvain-Jaccard algorithm (Shekhar et al., 2016). Integration of sc and snRNA-seq enabled identification of a total of 125 LAM^CORE^ cells but snRNA-seq had little effect on signature gene identification due to low gene detection sensitivity in the snRNA-seq LAM^CORE^cluster. We therefore combined data sets (LAM1, LAM3, LAM4sn) for further LAM cell sub-clustering, but excluded snRNA-seq data for analysis of differentially expressed genes in LAM^CORE^ cells.

### Identification of LAM^CORE^ cell subtypes

Mesenchymal cells in control human lungs were identified from scRNA-seq of seven female donor lung samples (**Figure S5**). Data from individual samples were preprocessed individually and integrated using the data integration function in Seurat 3 (Stuart et al., 2018). Cell clusters were identified using the Louvain algorithm implemented in Seurat 3 (Stuart et al., 2018). Cluster of cells with selective expression of lung mesenchymal markers were identified as Mesen^CONTROL^. Mesen^CONTROL^ cells from female donor lung samples were integrated with the LAM^CORE^ cells and Mesen^LAM^ (LAM associated mesenchymal) cells from scRNA-seq of the three LAM lungs using canonical correlation analysis in Seurat 2 (Butler et al., 2018), and clustered using the Louvain-Jaccard algorithm (Shekhar et al., 2016) to identify mesenchymal subtypes. Cluster specific differential expression was tested using the negative binomial test based method.

### Prediction of LAM^CORE^ cell origins

Pseudo-bulk samples were generated from scRNA-seq data as follows: for each gene, counts were summed across the cells of interest, normalized by total number of counts of all genes, and scaled by 100,000. Microarray data from the Human U133A/GNF1H Gene Atlas (Su et al., 2004) was downloaded from BioGPS (http://biogps.org/). RNA-seq data from Illumina’s Human BodyMap 2.0 was downloaded from Gene Expression Omnibus (GSE30611). It consists of RNA expression in 16 human tissue types. Expression data plus 1 were log2 transformed. The expression of genes (n=11,194) commonly detected in data from different platforms was used for batch correction using ComBat (Johnson et al., 2007). Principal component analysis on ComBat-corrected expression of 1,120 genes with highest coefficient of variation (10%) was performed for dimension reduction. The first 9 principal components were used for unbiased hierarchical clustering using Pvclust (Suzuki and Shimodaira, 2006) using Pearson’s correlation based distance, average linkage, and 1,000 bootstrap replications.

### LAM^CORE^ secretome

Secreted protein lists were obtained from the Human Protein Atlas (https://www.proteinatlas.org/humanproteome/cell/secreted+proteins) and Vertebrate Secretome Database (http://genomics.cicbiogune.es/VerSeDa). In the present study, “Refined Predictions Curated Set” (RPCS) from VerSeDa (Cortazar et al., 2017) and all secreted proteins from Human Protein Atlas were included for analysis. The Human Protein Atlas predicted a complete set of human secreted proteins (“secretome”) using three signal-peptide based prediction methods across 32 human tissues (Thul et al., 2017). VerSeDa combined secretome information from every vertebrate species available at the NCBI, University of California Santa Cruz (UCSC) and ENSEMBL databases, in addition to relevant model organisms’ proteomes (Cortazar et al., 2017). Secretome proteins are identified by the presence of a signal peptide, and not all are secreted into the extracellular space; some are retained in secretory pathway compartments, such as the endoplasmic reticulum or Golgi, or remain attached to the outer face of the cell membrane by a glycosylphosphatidylinositol anchor (Thul et al., 2017; Uhlen et al., 2015). Genes encoding secreted proteins were mapped to the LAM^CORE^ signature gene list (**Table S2**) and functional enrichment analysis was performed using ToppGene suite (Chen et al., 2009). A network diagram was generated using Cytoscape (Shannon et al., 2003) v3.7.1 for genes associated with the most enriched gene ontology biological processes and pathways.

### SOMAscan assay

The SOMAscan proteomic assay was performed as previously described (Gold et al., 2010). In brief, each of the 1,125 serum proteins measured in serum has its own targeted SOMAmer reagent, which is used as an affinity binding reagent and quantified on a custom Agilent hybridization chip. TSC, TSC-LAM and control samples were randomly assigned to plates within the each assay run along with a set of calibration and normalization samples. No identifying information was available to the laboratory technicians operating the assay. Intrarun normalization and interrun calibration were performed according to SOMAscan v3 assay data quality-control procedures. SOMAscan proteomic data are reported in relative fluorescence units (RFU), which were log-transformed before statistical analysis to reduce heteroscedasticity. Differentially expressed genes were defined using FDR of One-tailed Welch’s t-test *p* value <0.001 and fold change >=1.5.

### ELISA assays

Endostatin and IGFBP-4 serum concentrations were determined with an enzyme-linked immunosorbent assay (ELISA; R&D systems, Minneapolis) per manufacturer’s instructions.

### Histology and immunohistochemistry analysis and immunofluorescence microscopy

Tissues for histologic and immunohistochemical analysis were fixed in formalin, embedded in paraffin and sectioned. Tissue sections were stained with hematoxylin and eosin for histopathologic analysis. Immunohistochemistry was performed with the Roche Ventana BenchMark Ultra IHC/ISH System using the following antibodies: smooth muscle actin (IA4, Roche catalog# 760-2833), Melanosome (HMB-45, CONFIRM, Roche catalog# 790-4366), MART-1/melan A (A103, CONFIRM Roche catalog# 790-2990) and Podoplanin (D2-40, BioLegend catalog# 916604). A Complete list of antibodies used in the present study was shown in **Table S9**. Images were taken with a Nikon Eclipse 80i light microscope using a SPOT camera and SPOT Software 5.2. Alternatively, tissues were fixed overnight in 4% paraformaldehyde/PBS, equilibrated in 30% sucrose, and embedded in OCT or fixed in 10% formalin and embedded in paraffin. F7 micron frozen sections (0.7micron) or FFPP sections (10 micron) were used. Immunofluorescence staining was performed using methodologies previously described at https://research.cchmc.org/lungimage. Antibody information and working dilutions are indicated in **Table S9**. Confocal microscopic images were taken on instruments Nikon A1 LUNA inverted, Nikon A1R LUN-V inverted, Nikon A1R inverted, using Nikon elements software. Images were post -processed in either Nikon Elements or IMARIS BITPLANE. Brightfield images were acquired on a Zeiss AXIOIMAGER.A2 using AxioVision software. Sections (5μm) of paraffin embedded normal donor and LAM lungs were baked at 50°C overnight. Sections were deparaffinized, rehydrated in a series of 100% EtOH, 95% EtOH, 70% EtOH and cleared in phosphate buffered saline (PBS) for five minutes and treated with sodium citrate for antigen retrieval. Normal goat serum (4%) in PBS, 0.1% Triton-X (PBST) was used for blocking. Slides were incubated in primary antibodies (**Table S9**) diluted in blocking buffer for 12-24 hours at 4°C were washed in PBST. Secondary antibody (1:200) and DAPI (1mg/ml) were added for 1 hour at room temperature. Samples were washed in PBST and mounted with Prolong Gold (Thermo Fisher Scientific). Images were acquired on an inverted Nikon A1R confocal microscope. Maximum intensity projections of *Z*-stack images were generated using NIS-Elements software (Nikon).

### Quantitative RT-PCR

RNA from snap-frozen lung tissues was isolated using the RNeasy Mini Kit (Qiagen). Gene expression was quantified using One-Step qRT-PCR Kits (Invitrogen) in the Applied Biosystems Step One Plus Real-Time PCR System and normalized to β-actin (*ACTB*) RNA. Each target gene was tested in triplicate for each lung sample. RNA assay primers are listed in **Table S9**.

### Statistics

To identify differentially expressed genes in scRNA-seq and snRNA-seq analyses, statistical analyses were performed using a one-tailed binomial probability based test (Shekhar et al., 2016) and a modified negative binomial probability based test implemented in SINCERA pipeline (Guo et al., 2015; Yu et al., 2013). One-tailed Welch’s *t* test was used for statistical analysis of Aptamer proteomics data. Student *t* test was used for two groups, and one-way ANOVA test was used for multiple group comparison in ELISA assays. Real-time RT-PCR data represent mean ± SEM. Statistical analyses were performed using two-tailed Student *t* test when comparing two groups, and one way ANOVA test (Dunnett’s multiple comparisons test when comparing multiple groups with control group, Tukey’s multiple comparisons test when making multiple pairwise comparisons between different groups) for multiple group comparison. A *p* value less than 0.05 was considered significant.

## Acknowledgements

This research was supported by the LAM Foundation (LAM0138E01-19 to A.K.P. and Y.X.; LAM0131PB07-18 to A.K.P.). and the NHLBI (U01HL122642 to J.A.W., S.S.P., and Y.X.; U01HL134745 to J.A.W., Y.X., and M.G.; R01 HL131661 to AKP and YX, R01HL138481 to J.J.Y.; T32 HL007752 to J.A.W. and M.R, U01 U54HL127672 and U01HL131022 to FXM). The LAM Foundation and NDRI assisted with tissue collection. This study was also supported by services from the Pathology Research Core shared facility in the Cincinnati Children’s Research Foundation with specific acknowledgement of the assistance of Betsy DiPasquale; and supported by services from UC Pathology for providing normal uterine tissue, and St. Vinvent Women’s Hospital for LAM-uterine tissue, and Cincinnati Life Center for normal lung. We thank Andrew Wagner for data mining assistance.

## Author contributions

F.X.M, J.A.W., K.A.W-B., and Y.X. conceived and designed the experiments. M.G., P.S., Y.X. designed and performed LAM single cell analysis and associated integrative bioinformatics analyses and interpreted data. E.Y.Z., and J.J.Y. designed the validation experiments using qRT-PCR and immunofluorescent staining. A.K.P and M.R designed and analyzed LAM^CORE^ cells and subtypes spatial location and signature genes validation using immunofluorescent confocal imaging. K.G. and D.J.K. conducted mutation analysis. F.X.M designed and validated LAM secretome findings in LAM patient’s serum using Aptamer screen and ELISA. M.A., A.P., and S.S.P. conducted single-cell RNA-seq experiments and contributed to single cell data analysis. E.J.K. coordinated tissue collection and regulatory issues. K.A.W-B examined, processed and pathologically evaluated human tissues, selected human lung and uterus samples for single cell and single nuclei RNA-seq experiments, confirmed lung and established uterine pathologic diagnoses, and determined immunophenotype of LAM cells. M.G., Y.X., F.X.M., and J.A.W. wrote the manuscript. All authors contributed to the data interpretation, troubleshooting, manuscript writing and editing. All of the authors read and approved the final manuscript.

